# The RNA-Binding Protein SAM68 regulates cardiomyocyte differentiation by enhancing Gata4 translation

**DOI:** 10.1101/2022.01.11.475875

**Authors:** Alessandro Dasti, Maria Carla Antonelli, Magdalena Arnal Segura, Alexandros Armaos, Sarah Bonnin, Thomas Graf, Maria Paola Paronetto, Tian V Tian, Elias Bechara, Gian Gaetano Tartaglia

## Abstract

The signal transduction and activation of RNA (STAR) family is composed of RNA-binding proteins (RBPs) that play a central role in mammalian development. Nonetheless, the functions and modes of action that STAR proteins have in lineage specification are still poorly understood. Here, we characterized the role of STAR proteins SAM68 and QUAKING (QKI) in pluripotency and differentiation by performing their depletion through CRISPR-Cas9 in mouse embryonic stem cells (mESCs). Combining RNA-sequencing, ribosome profiling and advanced computational predictions, we found that both SAM68 and QKI regulate the mESCs self-renewal and are indispensable for cardiomyocyte differentiation. At the molecular level, we discovered that SAM68 and QKI antagonistically control the expression of cardiogenic factors. Our calculations indicated that SAM68, unlike QKI, binds the cardiogenic-specific transcription factor Gata4 in a region spanning nucleotides 500 to 1000 of the mRNA corresponding to part of the 5’ untranslated region and the first exon. We validated the predictions by electrophoretic mobility shift assay and RNA immunoprecipitation showing that SAM68 controls the translation of Gata4 during mESCs differentiation towards the cardiomyocyte lineage.

## Introduction

Embryonic stem cells (ESC) are a laboratory tool to study the early phases of mammalian development in addition to being promising in both therapy and disease modelling ^1,2^. The mechanisms underlying the self-renewal and pluripotency properties of the ESCs have been subject of intensive studies in recent years ^3–5^. Upon differentiation stimuli, the ESCs actively exit from the pluripotent state by disrupting the core transcriptional network maintained by master pluripotency transcription factors (TFs) and by expressing lineage-specific transcription factors that in turn activate gene expression programs leading to differentiation ^6^. In addition to TFs and and chromatin associated factors involved in transcription regulation, advances in - omics techniques such as RNA-seq have provided strong evidence that the post-transcriptional regulation is involved in pluripotency establishment, maintenance and lineage commitment ^7–9^. RNA Binding Proteins (RBPs) have been shown to be major players regulating the different steps of the RNA metabolism in this process ^10–14^. For example, the RBPs FOX2, SON, SFRS2, MYC, GCN5, ZCCHC24, and RBM47 facilitate pluripotency-specific alternative splicing (AS) of their target genes ^15–18^. In contrast, MBNL1, MBNL2, and SFRS11 promote differentiation-specific AS patterns for a large number of splicing events. Furthermore, RBM24 enhances cardiac-specific splicing favoring the cardiomyocyte differentiation of ESCs ^19^.

Another family of RBPs, STAR (Signal Transduction and Activation of RNA), belongs to the hnRNP K-homology (KH) domain family of proteins. It is composed of five members including KHDRBS1, also known as SAM68, and QUAKING (QKI) and is involved in mammalian development. Both SAM68 and QKI are regulators of AS, translation and stability of RNAs. The Sam68^-/-^ mouse model shows complete infertility in males and impedes the onset of the mammary gland tumor in females ^20,21^. Indeed, SAM68 plays a critical role during male germ cells development by enhancing the translation of RNAs that regulate spermatogenesis ^22,23^. Moreover, SAM68 is important in fine tuning the switch between self-renewal and differentiation of the neuronal progenitor cell ^24^. Although most Sam68^-/-^ mice are viable, many die at birth of unknown causes Richard et al., 2005). In contrast, the QKI^-/-^ mouse model is not viable due to severe cardiovascular defects suggesting an important role of this protein during cardiac tissue development ^25–27^. Indeed, QKI was recently shown to be critical for cardiac myofibrillogenesis and contractile function ^28^. Nonetheless, the mechanisms underlying the involvement of Sam68 and QKI in the early stage of mammalian development are still largely unknown.

In this study, we investigated the role of SAM68 and QKI in the lineage specification of mouse embryonic stem cells (mESCs). We showed that the depletion of either protein by CRISPR-Cas9 has a distinct effect on ESCs self-renewal and proliferation. Using RNA sequencing we identified several pathways involved in heart development to be mis-regulated in Sam68^-/-^ or QKI^-/-^ cells that fail to form bona fide cardiomyocytes. Combining computational predictions and ribosome profiling techniques we reveal hundreds of transcripts, including many implicated in proper cardiomyocyte differentiation, which are directly bound and/or whose expression is modulated by SAM68 or QKI. We specifically showed that SAM68 bind directly to the transcription factor Gata4 mRNA and regulate its translation. Our findings revealed a key circuit of posttranscriptional gene regulation operated by SAM68 and QKI that can be disrupted during cardiomyocyte differentiation.

## Results

### SAM68 and QKI positively regulate cell cycle, self-renewal in ESCs and cardiomyocyte differentiation

To study the role of SAM68 and QKI in early development, we engineered mouse embryonic stem cells (mESC) (E14TG2a) depleted of either Sam68 or QKI using the CRISPR-Cas9 technology (Figure 1A; Materials and Methods; Supplementary Material). Because the sgRNA were designed to target the first exon of the genes, this resulted in a complete abolishment of the proteins compared with wild type cells (Figure 1A). Importantly, the absence of any of the two proteins did not significantly affect the expression level of the other (Figure 1A). Knowing that QKI was involved in cardiac myofibrillogenesis, we performed in vitro cardiomyocytes differentiation and quantified the spontaneous beating foci after 8 days of induction of differentiation. In line with previous evidence, we found a significant reduction of beating foci in absence of Qki, and, unexpectedly, the same effect was observed upon KO of Sam68 (Figure 1B; Materials and Methods; Supplementary Material), indicating that like QKI, SAM68 has an important role in this specific differentiative path. In addition, the foci generated in the absence of Sam68 beat faster compared to the WT counterparts (Figure 1C). However, the very reduced number of Qki^-/-^ beating foci and their altered shape did not allow us further quantification. To better understand the origins of this phenotype, we characterized the mESC KO models. As the wild type mESC (WT), Sam68^-/-^ and Qki^i-/-^ knockouts mESC (KOs) showed normal domed-shape colony morphology with defined borders and differentiated colonies with spindle-shape were absent (Figure S1B).

**Figure 1.**
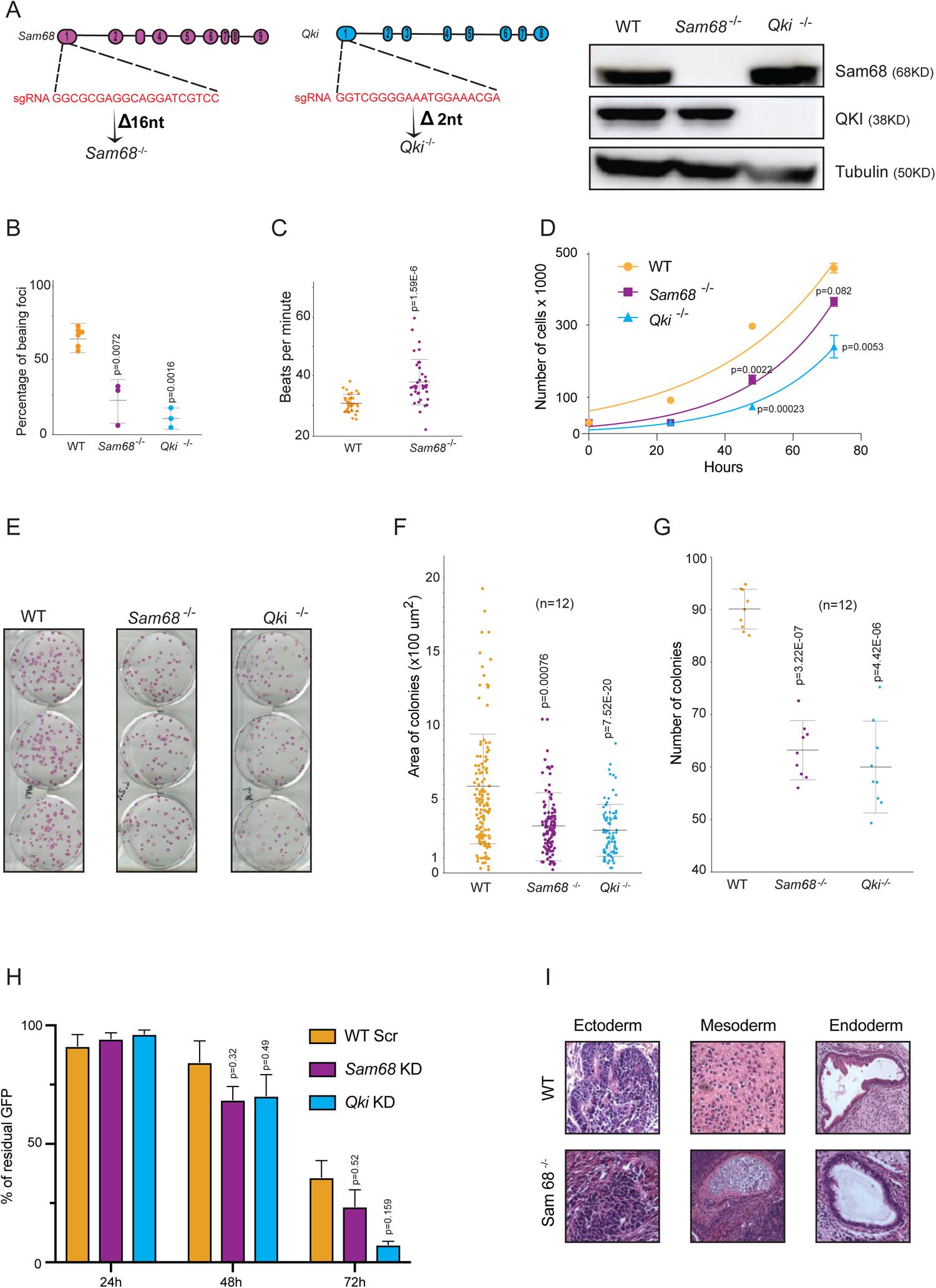
SAM68 and QKI positively regulate cell cycle and self-renewal in ESCs. a) CRISPR-Cas9 design strategy targeting the first exon of either Sam68 or QKI genomic locus in order to generate E14 mESC KO lines. The induced frame shift in both cases is shown in the Sanger sequencing of the respective cell lines. On the right: Western blot validating the absence of either protein. b) Reduction of beating foci in the absence of either QKI or SAM68 compared to WT. c) The foci generated in the absence of Sam68 beat faster compared to their WT counterparts. d) Proliferation assay of WT, Sam68^-/-^ and Qki^-/-^ mESCs. Each time point is the average of three independent biological replicas. The p-values obtained by student t-test are indicated for each significant point. e) Clonogenic assay pictures of WT, Sam68^-/-^ and Qki^-/-^ mESCs. f) Area of the colonies in μm^2^. Each column represents the results obtained by performing the experiment in three biological replicas, p-values for Sam68 ^-/-^ and QKI^-/-^ are indicated. g) Total number of colonies obtained after plating 200 cells per well. Results are represented for the three biological triplicates h) Exit from pluripotency. The bar plot shows the percentage of residual GFP (pluripotent cells) at different time points. i) Teratoma assay indicating the presence of terminally differentiated cells derived from ectoderm, mesoderm and endoderm

To study whether SAM68 and QKI were involved in pluripotent stem cell self-renewal, we first synchronized WT and KO cells with 400nM of nocodazole for 14-16 hours. Cell proliferation was monitored by daily cell counting through 72 hours (Materials and Methods; Supplementary Material). We found that depletion of either protein resulted in a significant decrease in the proliferation rate (Figure 1D). In agreement with this finding, the KO cells formed smaller colonies with reduced colony numbers compared to WT mESCs, in the clonogenic assay (Figures 1E, 1F and 1G). Taken together, our results indicate that both Sam68 and QKI positively sustained the self-renewal of the mESCs. This observation is consistent with a previous study associating the role of SAM68 in the self-renewal of neuronal progenitor cells (NPCs) ^24^. As mESC self-renewal is sustained by two inhibitors LIF and BMP, we removed them to provoke spontaneous differentiation. We therefore used Rex1-d2GFP reporter cells to assess whether SAM68 and QKI were involved in this process ^29^. We first inhibited the expression of Sam68 and QKI in Rex1-d2GFP cells by infecting cells with lentiviral particles harboring short-hairpin RNA (shRNA) constructs against Sam68 and Qki (KD) (Figure S1C; Materials and Methods).

The dynamics of pluripotency exit of these cells was measured by FACS after 24hrs, 48hrs and 72hrs after removal of the two inhibitors and growing the cells in differentiation-permissive medium. As shown in Figure 1H, there was no significant difference between control and KD cells, suggesting that none of the proteins is involved in this process (Figure 1H). Next, to further evaluate the involvement of both proteins in early lineage specification in vivo, we implanted subcutaneously the KO mESCs and their WT counterpart into Nude mice to form teratomas. Interestingly, teratomas formed by either WT or KO mESCs derived cells showed cells of all three germ layers (Figure 1I). The Sam68^-/-^, but not Qki^-/-^ teratomas showed an increased cell from mesodermal origin (Figure S1D), suggesting that Sam68 protein may be involved in the regulation of mesoderm differentiation.

## Is the cardiomyocyte phenotype due to transcriptional or post-transcriptional regulation by SAM68 and QKI?

To assess the roles of SAM68 and QKI during early mammalian development, we allowed WT and Sam68^-/-^ or Qki^-/-^ ESCs to spontaneously differentiate as tissue-like spheroid in a suspension culture forming embryoid body (EB) for ten days (Materials and Methods; Supplementary Material). This assay recapitulated the early stages of mammalian development with the formation of a complex three-dimensional structure made of cells and extracellular matrix that drove the differentiation of the cells towards the three germ layers and their derivatives. SAM68 and QKI protein levels were checked throughout the differentiation in wild type and knockout cells (Figure S2A). Furthermore, RNA was isolated and used to perform total RNA-sequencing in three biological replicates in different time points (Day0, Day3, Day10; Materials and Methods; Supplementary Material).

We first performed an unbiased principal component analysis (PCA) that allowed us to detect significant and consistent changes in gene expression in Sam68^-/-^ EBs (Figure S2B), whereas Qki^-/-^ EBs showed a more similar pattern to its WT counterparts (Figure S2C). Although the expression of all the classes of coding and non-coding RNAs was affected at each time-point of differentiation (Figure 2A), the extent of the expression deregulation in Sam68^-/-^ EBs was more prominent compared to Qki^-/-^ EBs (Figure 2B, Figure S2D) suggesting a broader role of Sam68 in regulating all the RNA types during ESCs differentiation. Interestingly, the overlap between differentially expressed RNAs for the two proteins in each time-point was rather limited (Figure 2C), revealing distinct regulatory functions of these factors in the differentiation towards various lineages. Furthermore, the percentage of regulated transcripts in opposite directions of the overlapping cluster increased from 28% to 62% in Day0 and Day10 respectively (Figure 2C; Supplementary Material).

**Figure 2.**
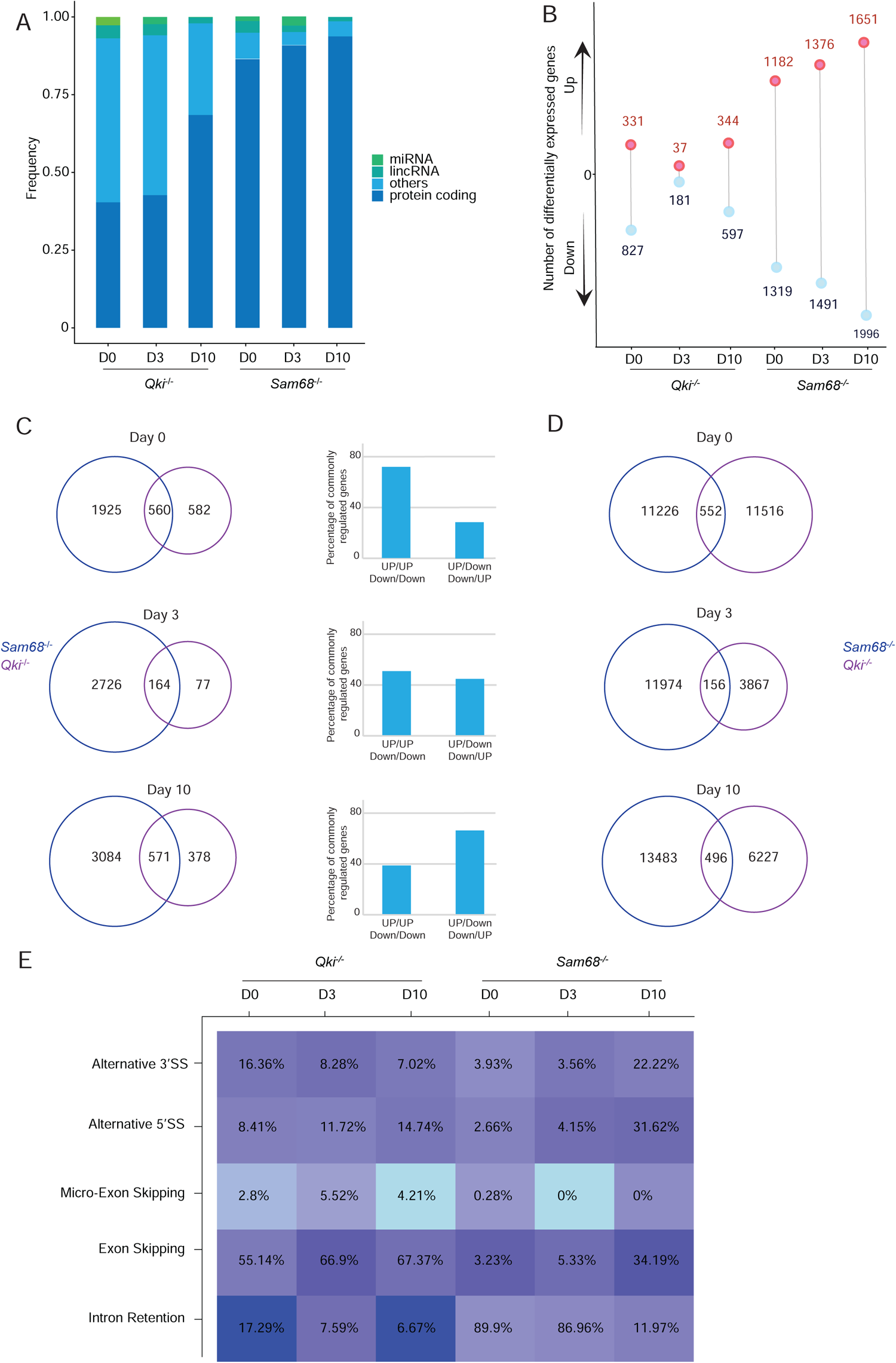
Transcriptional and posttranscriptional landscape driven by SAM68 and QKI during ESCs differentiation. a) Different classes of differentially expressed RNAs in either Sam68-/-or Qki-/-mESCs at day 0, day 3 and day 10 of EBs differentiation. Only RNAs with ± 1,5 fold change and an adjusted p value<0,01 are represented. b) Numbers of differentially expressed protein coding RNAs at each time point for each KO cell line c) Deregulated coding RNAs at each time point of both Sam68^-/-^ and QKI^-/-^ cells. Their intersection together with the percentage of commonly regulated in the same direction (up/up or down/down) and in opposite directions (up/down or down/up) for each protein. d)Deregulated alternative splicing events in the two cell lines at day 0, day 3, day 10 of EBs differentiation. The intersection of the different sets is less than 5% e) Heatmap showing the different types of aberrant AS events in the absence of either QKI or SAM68 during EBs differentiation of mESCs. Only events with a DPSI>+25% or <-25% and an adjusted p value<0,01 are considered.

At the post-transcriptional level, both SAM68 and QKI were previously described to regulate the AS in different developmental processes ^30–33^. The global changes of the AS in Sam68^-/-^ and QKI^-/-^ EBs compared to WT EBs were therefore measured. We found that despite the large number of deregulated events in each cell line at each time point, the overlap between the two KO lines was less than 5%, indicating divergent roles of the two proteins in the AS regulation in pluripotent mESCs and during their differentiation (Figure 2D). This pattern of regulation has been depicted for members of the SR RBPs family ^34,35^ and the RBM family ^36^ Surprisingly, in the absence of SAM68 the majority of the altered AS events belonged to the category of intron retention, in line with what was previously described for specific SAM68 targets ^37,38^ whereas cassette exon and alternative 3’ or 5’ splice site choices appeared to prevail in QKI^-/-^ (Figure 2E) suggesting distinct mechanism through which SAM68 and QKI regulated the AS.

## Do SAM68 and QKI regulate cardiac-specific RNAs?

We performed Gene Ontology (GO) analysis on the differentially expressed genes at day 10 of EBs. In addition to genes involved in neurogenesis and vasculature development as previously described ^30,33,39^ (Figures S3A and S3B), an overrepresentation of cardiac related terms was observed implicating both SAM68 and QKI in cardiac development. Surprisingly, while the majority of cardiac-related RNAs were upregulated upon the depletion of SAM68, a complete opposite pattern was observed in Qki^-/-^ EBs (Figure 3A). In the absence of SAM68, we observed significant upregulation of Gata4, Nkx2-5 and Tbx18 and downregulation of Isl1 in D10 EBs (Figure 3B). This general imbalance of cardiac progenitor specific TFs ^40^ was likely to be the reason underlying the general upregulation of transcripts encoding structural (Tnnt2, Myl2, Myl4, Actc1) ^41,42^ and functional proteins (Cacnb2 ^43^, Ryr2) of the cardiac cells (Figure 3B). In QKI^-/-^ we found a downregulation of the expression of cardiomyocyte structural proteins (e.g., Actn2, Mybphl, Nebl, Myh6, Actc1, Tmod1, Tnni3, Tnnc1, Tnnt3) ^19,44–49^, functional proteins, (e.g., ion channels Ryr2 and Scn5A)^50– 52^, and cardiac-specific splicing regulators (Rbm20 and Rbm24)^19,44^ that were confirmed by RT-PCR (Figure 3C). This downregulation can be partially explained by the significant downregulation of master cardiac TFs Gata4, Gata6 and Mef2c, detected in the earlier stages of differentiation in the Qki^-/-^ cells (Figure 3D). Indeed, these TFs are responsible for the expression of many of the genes that are downregulated at day10 of EBs differentiation ^53,54^. The deregulation at the expression level came along with an altered pattern of AS in both Sam68^-/-^ and Qki^-/-^ cell lines. More specifically, the absence of QKI affected the correct inclusion of several cassette exons of structural and functional cardiac components (Figure S3C) whereas the absence of SAM68 led to the aberrant splicing of different exons of the calcium channel subunit, Cacna1c, which is highly important for the excitability of the cardiomyocytes (Figure S3D)^43^. Although the phenotype that we have observed in Sam68^-/-^ was identical to the one reported using Gata4^-/-^ mESCs in vitro ^55^, this could be related either to the aberrant expression of Ryr2 or the altered AS of Cacna1c (or a combination of both). To sum up, our data indicate that both SAM68 and QKI are important factors in the regulation of the cardiomyocyte differentiation of mESCs through the control of cardiac-related RNAs expression, AS and their subsequent translation.

**Figure 3.**
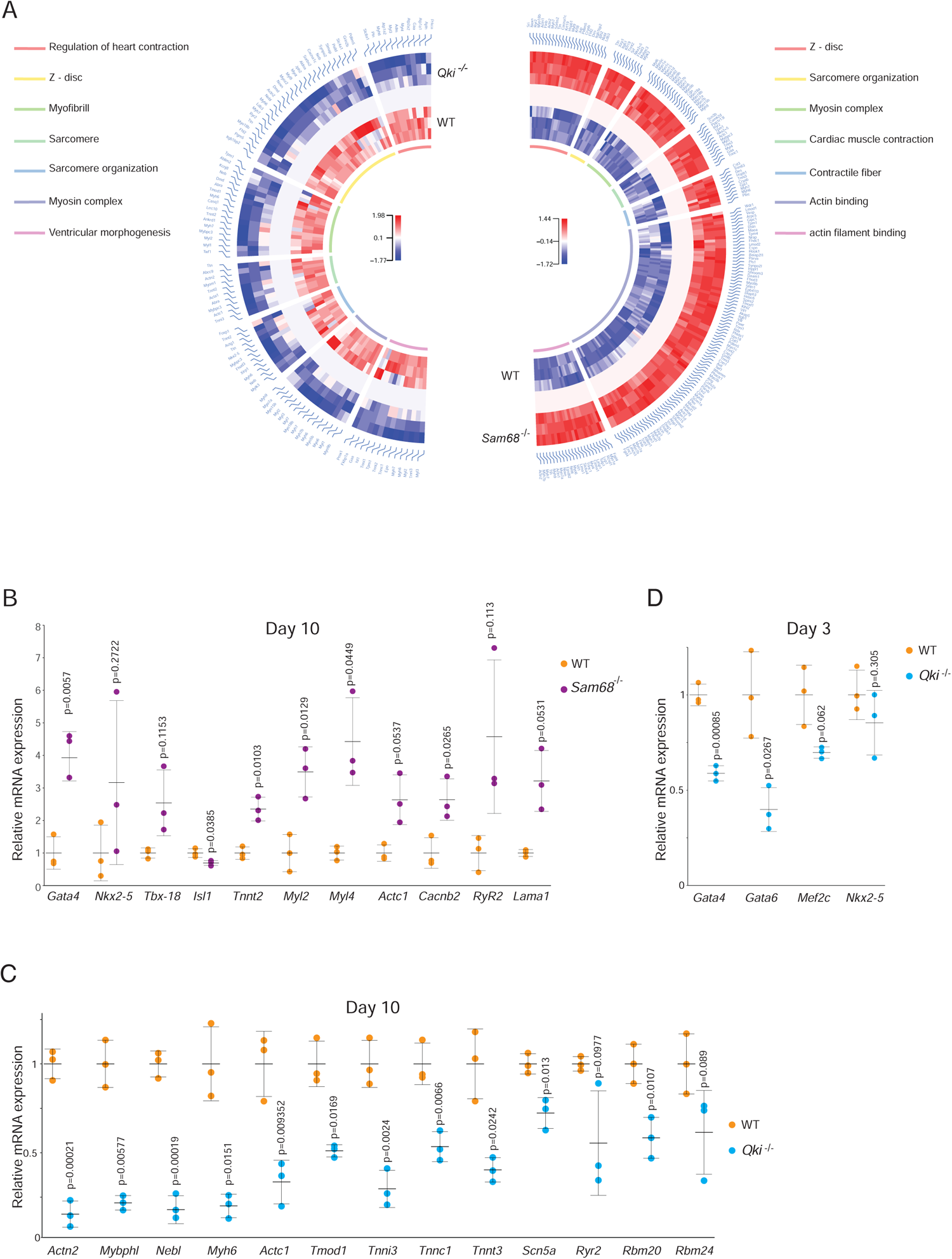
SAM68 and QKI regulate cardiac-specific RNAs and cardiomyocyte differentiation. a) Circle plot of selected GO gene-sets related with cardiac development at day 10 of EBs differentiation. RNAs belonging to this category are mainly downregulated for QKI^-/-^ cells (on the left) and upregulated for Sam68^-/-^ cells (on the right). GO classes are indicated in the respective color legends b) RNA-sequencing validation via real-time PCR of the altered expression of cardiac-related mRNAs in Sam68^-/-^ cells at day 10 of EBs differentiation. P-values are indicated for each validated transcript. c) RNA-sequencing validation of cardiac related transcripts in QKI^-/-^ EBs at day 10 of differentiation via real-time PCR shows severe downregulation of these RNAs. d)Real-time PCR shows a significant downregulation of Gata4 and other cardiogenic transcription factors in QKI^-/-^ ells at day 3 of EBs differentiation.

## Sam68 directly binds GATA4 mRNAs and regulates its translation

The upregulation of Gata4 mRNA in Sam68^-/-^ could be accompanied by more significant cardiomyocyte differentiation ^56^, however the observed phenotype reflected its absence ^55^. This result could be explained considering Gata4 mRNA post-transcriptional regulation. Indeed, both SAM68 and QKI were previously described to regulate translation ^23^. To assess the global variation in translation in Sam68^-/-^ and Qki^-/-^, ribosome profiling was performed on day 10 of EBs (Materials and Methods; Supplementary Material). This technique snapshots all the actively translated mRNAs ^57^. The majority of the obtained clusters aligned to coding sequences indicating high-quality library preparation (Figure S4A). Cardiac transcripts whose translation was significantly affected were found in Sam68^-/-^ but not in Qki^-/-^ suggesting that SAM68 was involved in both post transcription and translation process of cardiac factors, whereas the role of QKI was translation independent (Figures 4A and S4B). SAM68 repressed the translation of ion channels (Kcnj8)^58^, structural proteins (Myh6) and TFs (Nkx2-5, Osr1 ^59^). Conversely, it enhanced the translation of the mesoderm specifier transcription factor Sox17 ^60^, Ihh, a mediator of the Indian Hedgehog signaling pathway indispensable for the embryonic development of the heart ^61^ and most importantly of Gata4 (Figure 4A) which was further confirmed by western blot and immunofluorescence (Figures 4B, 4C and S4C).

**Figure 4.**
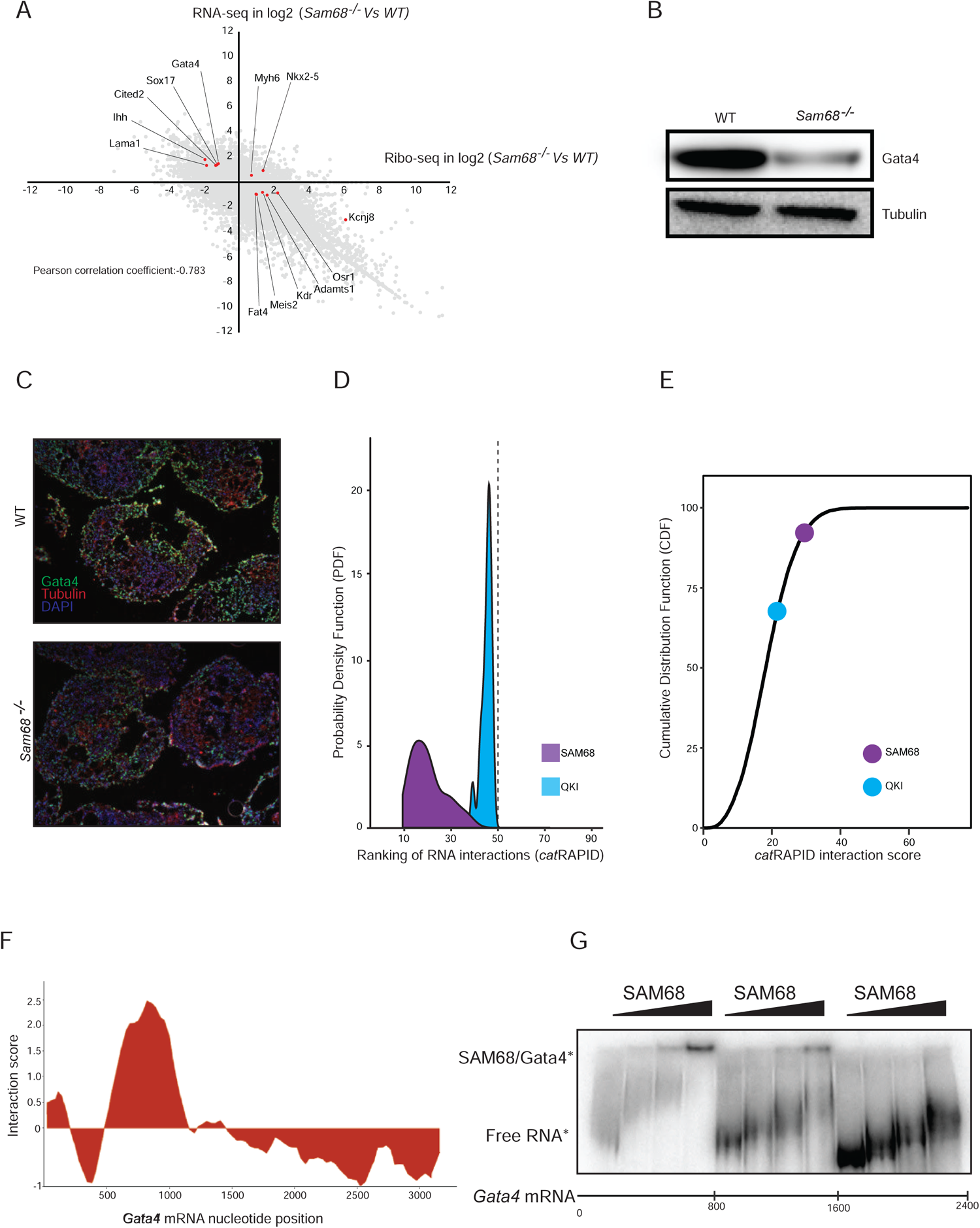
SAM68 directly binds Gata4 mRNAs and regulates its translation. a) Transcription-translation correlation plot in Sam68^-/-^ EBs at day10 of differentiation. Cardiac-related RNAs whose translation are significantly affected are highlighted b) Western blot showing a dramatic reduction of GATA4 protein levels in Sam68^-/-EBs^ c) Immunofluorescence showing reduced GATA4 (in green) levels in the absence of Sam68. d)Plot showing the interaction propensity of either SAM68 or QKI to cardiac-related transcripts whose translation is affected in absence of Sam68 in differentiating EBs compared to a pool of 1799 RBPs. e) Plot showing the cumulative distribution function (CDF) of the catRAPID score calculated for QKI and SAM68 to the mRNA mouse transcriptome with RNAs ranging from 3000 to 4000 nucleotides. The dots represent the binding propensity of either QKI or Sam68 to Gata4 RNA. f)Interaction propensity between SAM68 protein and Gata4 RNA as predicted by catRAPID g) EMSA shows that SAM68 binds to the first part of Gata4 RNA as demonstrated by the shift of the SAM68-Gata4 RNA complex

To unravel whether SAM68 regulated Gata4 translation through direct binding to its mRNA, we first calculated the interaction propensity of the affected cardiac mRNAs in Sam68 and QKI KO EBs using catRAPID ^62,63^ (Materials and Methods) The catRAPID algorithm estimates the potential of a protein RNA pair through van der Waals, hydrogen bonding and secondary structure propensities predicted from their sequences. Comparison with 1799 high-confidenceRBPs (Materials and Methods) reveals that SAM68 has a higher interaction propensity compared to QKI as it falls in the top 20% of all RBPs in the probability distribution function (PDF; Figure 4D). In addition, according to the cumulative distribution Function (CDF), SAM68 shows an interaction to Gata4 mRNA around 90% higher compared to the rest of the transcriptome (Figure 4E). This interaction is predicted to occur in a region spanning nucleotides 500 to 1000 of the Gata4 nascent mRNA corresponding to part of the 5’ untranslated region (UTR) and the first exon of the transcript (Figure 4F). Importantly, we further validated this finding by electrophoretic mobility shift assay (EMSA) (Figure 4G) and RNA immunoprecipitation (RIP) assay (Figure S4D; Materials and Methods)). To conclude, during mESCs differentiation towards the cardiomyocyte lineage, SAM68, unlike QKI, binds directly and controls the translation of the cardiogenic transcription factor Gata4.

## Discussion

Here we described how two members from the same RNA Binding Protein family could have overlapping, but also distinguishable roles in both embryonic stem cells pluripotency and differentiation. Our results indicate that both SAM68 and QKI positively regulate mESCs proliferation, self-renewal and are indispensable for proper differentiation towards the cardiomyocyte lineage by regulating expression, alternative splicing and translation of transcription factors, structural and functional cardiac-specific proteins. In particular, we show that Gata4 is a direct target of SAM68 which regulates its physiological protein levels.

### SAM68 and QKI in self-renewal and proliferation

Clonogenic assay demonstrated that both SAM68 and QKI are involved in proliferation and self-renewal of mouse embryonic stem cells. These proteins have been reported to regulate cell proliferation ^27,64^ and our findings go in line with previously reported prolonged G2-M phase in Sam68 depleted chicken fibroblasts ^65^ and an altered cell proliferation in absence of QKI in colon cancer cells ^66^. One of the molecular mechanisms orchestrating the reduced self-renewal capacities of both Qki^-/-^ and Sam68^-/-^ mESCs could be attributed to the decreased levels of Wnt3 and its direct downstream target Lef1, known to promote self-renewal ^67^. Indeed, we observed the downregulation of both Wnt3 in Sam68 deficient cells and Lef1 at the transcription and translation levels in Qki^-/-^ and Sam68^-/-^ respectively. On the other hand, mESCs use glycolysis as a main source of ATP production and switch to oxidative phosphorylation upon differentiation ^68^. This metabolic switch mechanism was shown to connect SAM68 to self-renewal in neuronal progenitor cells ^24^. In addition, our results demonstrate that the non-coding RNA, Lncenc1, recently shown to be important for the expression of glycolysis-associated genes ^69^ to be consistently downregulated in both Qki^-/-^ and Sam68^-/-^ cells.

### Role in cardiac differentiation and role of other RBPs

RNA-sequencing at both early and late stages of embryoid bodies (EBs) differentiation in SAM68 and QKI depleted cells demonstrated their implication in the nervous tissue development as shown ^24,27^. Strikingly, we also revealed novel pathways that were not previously described. Among these, we found terms containing genes encoding important functional and structural cardiomyocytes-specific proteins that are regulated in opposite directions in Sam68 and QKI KO cells. This suggests that not only both proteins are involved in the development of the heart but that they probably exert different functions in this biological context. This might not be surprising as it has been shown that RBPs belonging to the same family can exert antagonistic functions in the same process ^35,36^. In detail, QKI acts early on the core TFs network at day 3 of EBs differentiation. Although at this stage these TFs are lowly expressed ^70^, we observed a dramatic downregulation of Gata4 and Gata6 mRNAs. This downregulation was further extended to several genes that codify for structural, functional and cardiac-specific splicing regulators (Rbm24 and Rbm20) at day 10. The latter could be, at least partially, the reason underlying the aberrant alternative splicing landscape occurring in the cardiac specific transcripts ^19,71^. These extensive defects in the very early stages of differentiation, might be the reason underlying the lack of beating foci formation as well as the lethality of the QKI^-/-^ embryos. Conversely, in the case of Sam68^-/-^ cells, the majority of the defects were observed at D10 of EBs differentiation. In fact, we detected a dramatic upregulation of the mRNA levels of cardiac transcription factors (Gata4, Nkx2-5 and Tbx18) that form a cascade that drives precursor cells towards a cardiomyocyte cell identity ^72,73^. In addition, Tbx18 is known to be crucial for the formation of the sinoatrial node cells that initiate the electric impulse and stimulate the contraction of the organ ^40^. To the same extent, the depletion of Sam68 caused aberrant upregulation of structural proteins (Tnnt2 and Actc1) and the ion channels (RyR2 and Cacnb2) important for the cardiomyocyte depolarization and their consequent contractions ^74,75^. Moreover, Cacna1c, another important ion channel subunit ^76^, showed altered pattern of several cassette exons inclusions in absence of Sam68. All together, these data can explain the abnormal beating activity that we observed upon Sam68 depletion.

### SAM68 and Gata4

Pre-mRNA splicing, localization and stability were shown to be the molecular mechanisms through which QKI regulates neuronal development ^77–79^. Our results show that, unlike QKI, Sam68 can exert its role in cardiomyocyte differentiation through translation regulation as well. SAM68 as translation enhancer was previously described in male germ cells differentiation ^22,23^. Here we showed, for the first time, that the translation regulation mechanism through SAM68 is conserved in the development of the cardiac tissue. Nevertheless, our results presented a puzzling case where Gata4 showed sharp up and downregulation of its mRNA and protein levels respectively in the absence of SAM68. Moreover, we have shown that Sam68 binds directly to Gata4 mRNA and acts as an enhancer of its translation. Two possible hypotheses can be at the basis of this mis-regulation, the nuclear export and the transcription-translation feedback loop. One of the known functions of SAM68 is the nuclear export of the HIV transcripts into the cytosol of the infected cells in order to be translated ^80^. By a similar mechanism and in normal conditions, SAM68 would enhance the nuclear export of Gata4 mRNA consequently allowing its proper translation. Conversely, in the absence of Sam68, the majority Gata4 mRNA might be retained in the nucleus, thus being less available to actively translating ribosomes leading to low protein production. In parallel, a positive transcription-translation feedback loop previously described for circadian oscillators ^81^ can also explain this puzzling scenario: In the absence of SAM68, Gata4 mRNA is poorly translated, this induces an increase in Gata4 mRNA transcription by either a transcription enhancer cofactor or through direct binding of GATA4 to its own promoter ^82^. Furthermore, the translation of Gata4 is crucial for the proper spermatocyte differentiation ^83^ which is highly compromised in Sam68^-/-^ mice ^22,23^. To date, this is the first evidence of post-transcriptional regulation of Gata4 mRNA by a specific RNA binding protein. Consequently, this can shed more light on the understanding of the Sam68^-/-^ infertility phenotype through Gata4 regulation. To sum up, Sam68 and QKI regulate a circuit of genes involved in the development of the cardiovascular system that could be the cause of the embryonic lethality in QKI^-/-^ and the high perinatal mortality of Sam68^-/-^ mice.

## MATERIALS AND METHODS

### CRISPR/Cas9-mediated genome editing

Genome-editing was carried out by transfecting E14 cells with the bicistronic plasmid pSpCas9 PX459 (1μg per well in a 6-well plate) encoding for both the Cas9 and the sgRNAs targeting the first exon of either Sam68 or QKI. The transfected cells were then selected with puromycin and FACS sorted at clonal density in a 96 multi-well plate. Clones that were successfully edited were then Sanger-sequenced and the absence of the protein product verified by Western Blot. See Table S1 for gRNA sequences.

### Proliferation assay

To assess the proliferation rate, 7,5×10^4^ cells were plated in a gelatin pre-coated 12 multi-well plate. Alive cells were then counted daily by Trypan Blue staining and quantified with the Countess™ II automated cell counter. Each well was counted at least twice. The graph in Figure 1B represents the average and the standard deviation of 3 independent experiments.

### Clonogenic assay

For this assay, 200 cells were plated on a pre-coated 6 well plate and cultured with the E14 maintenance medium supplemented with LIF and 15% FBS. The cells were then left in culture for 10 days and the medium was changed every other day. At day 10 the colonies were stained with the Alkaline Phosphatase Staining Kit II (00-0055, Stemgent) according to the manufacturer’s instructions. The colonies were then counted and the area of each colony quantified using Image J software. The dot plot in Figure 1D represents the average of 3 independent replicas.

### Exit from pluripotency assay

The Rex1-GFP KD and scramble cell lines were used in order to assess the exit from pluripotency and the commitment of these cells. Briefly, the cells were allowed to differentiate by removing both LIF and the 2 inhibitors. Cells were then detached with the ACCUTASE^®^ detachment solution and resuspended in ice-cold PBS supplemented with DAPI at a concentration of 0.1μg/mL in order to detect the fraction of death cells. Cells were analyzed at the BD LSR II cytometer. We first gated for forward and side scatter, we excluded the doublets and the death cells from the final analysis and we plotted the amount of residual GFP expression after 24, 48 and 72h after LIF and 2i removal. The bar plot in Figure 1F represents the average of 3 independent replicas.

### Teratoma assay

All the experimental protocols were performed in accordance with the recommendations for the proper care and use of laboratory animals [local (law 32/2007); European (EU directive n° 86/609, EU decree 2001-486) regulations, and the Standards for Use of Laboratory Animals n° A5388-01 (NIH)] and were approved by the local ethical committee (CEEA-PRBB). For Teratoma formation, 100μl of solution containing 500.000 mouse ES cells and 1:15 matrigel were injected into both flank sites of SCID BEIGE mice. The teratoma formation was stopped after 3 weeks of development. The resulting tissue was washed with PBS, fixed with 4%PFA, paraffin embedded, sectioned and stained with haematoxylin/eosin. Sections were evaluated and the presence of ectodermal, mesodermal and endodermal-derived tissue was quantified.

### RNA-sequencing

In order to perform the RNA-sequencing, total RNA was extracted with the Maxwell 16 LEV simplyRNA Cells Kit (AS1270, Promega) according to manufacturer’s instructions from WT, Sam68^-/-^ and Qki ^-/-^ EBs in three biological replicates. DNA-free RNA was then quantified using the Nanodrop and quality checked with the Bioanalizer. rRNA depletion and library preparation was done with the TruSeq stranded total RNA Library prer Human/Mouse/Rat (20020596, Illumina). The sequencing was performed using 2×125bp paired-ends reads on a HiSeq 2500 sequencer with HiSeq v4 chemistry. The raw reads were de-multiplexed, by the bcl2fastq Illumina software. The Illumina Universal adaptor was trimmed off the raw reads with Skewer (version 0.2.2), option “-x AGATCGGAAGAG”. The quality of both raw and trimmed reads was assessed with the FastQC tool. Trimmed reads were aligned to the Mus musculus genome (mm10/GRCm38) with the STAR RNA-seq mapper (version 2.5.3a) using option “--quantMode GeneCounts” and the M14 version of the Gencode annotation in order to obtain raw counts per gene and per sample (the 4^th^ column of the resulting counts file that corresponds to a “reverse stranded” protocol was kept). Bam files were converted to bigwig format with Samtools, so as to be loaded into the UCSC genome browser. The DESeq2 R/Bioconductor package was used to assess differential expression at gene level between experimental conditions. A gene was considered differentially expressed if the corresponding FDR-adjusted p-value was < 0.01 and the absolute fold change in log2 scale was > 0.6. Last, we used vast-tools (version 2.0.0) in order to detect alternative splicing events. MATT was used to filter vast-tools combined output. In this case, a dPSI threshold of 25 (described as stringent by the authors) was applied to filter significant results.

### Computational analysis of the binding propensity

The catRAPID algorithm ^62^ estimates the binding potential through van der Waals, hydrogen bonding and secondary structure propensities of both protein and RNA sequences allowing identification of binding partners with high confidence. As reported in a recent analysis of about half a million of experimentally validated interactions, the algorithm is able to separate interacting vs non-interacting pairs with an area under the ROC curve of 0.78 (with False Discovery Rate FDR significantly below 0.25 when the Z-score values are > 2) ^84^.

The specificity of the interaction of SAM68 or QKI for transcripts whose translation is affected in absence of Sam68 in differentiating EBs (Figure 4D) was calculated by considering the binding propensity ^62^ multiplied by the RNA binding ability ^85^, as implemented in catRAPID omics v 2.0 ^63^. The background distribution employed consisted of 1799 RBPs retrieved from literature ^86^.

Similarly, the interaction of SAM68 or QKI with Gata4 (Figure 4E), was calculated by considering the binding propensity ^62^ multiplied by the RNA binding ability ^85^, as implemented in catRAPID omics v 2.0 ^63^. To avoid biases arising from i) RNA type and ii) RNA length, we restricted the analysis of mouse transcriptome to protein coding transcripts that have length in the range (3000 to 4000 nts), i.e. similar to ENSMUST00000067417 (Gata4) which is 3400 nucleotides. The interaction binding profile (Figure 4F) was computed using the catRAPID fragment module ^87^.

### Electromobility shift assay

Electrophoretic mobility shift assays were carried out using 10 fmol of radiolabeled probes incubated on ice for 15 min with His-RBM_Nter_ in presence of 200 ng/μL yeast tRNA. Mixtures were then loaded on native 5% polyacrylamide gels, and migration was carried out in 0.5%TBE at 150V for 2 hours, followed by gel drying and Phosphorimager analyses.

### QUANTIFICATION AND STATISTICAL ANALYSIS

For each information about the statistical analysis, the meaning of the error bars and the number of replicates for each experiment please refer directly to the figure legends of each experiment.

## DATA AND CODE AVAILABILITY

## Data and code availability statement

The datasets and code generated during this study are available as detailed in the Key Resources Table.

## Data availability

All the data of the RNA-sequencing and the Ribosome Profiling are available at GEO with accession number GSE153800 and GSE153799 respectively

## Computer code

The code used for analyzing and processing the PALM data can be found at https://gitlab.com/anders.sejr.hansen/palm_pipeline.

Please see Key Resources Table for a full list of all the codes and softwares and where to find them.

## Acknowledgements

We thank all members of G.G.T. group and especially Dr. Elsa Zacco. We are grateful to Dr. Rosario Avolio, Dr. Davide De Pietri Tonelli and Prof. Stefano Gustincich for illuminating discussions. The research leading to these results was supported by European Research Council [ASTRA n. 855923 to GGT] and H2020 [INFORE n. 825080 to GGT] projects.

