## Supplementary Materials for "The RNA-Binding Protein SAM68 regulates cardiomyocyte differentiation by enhancing Gata4 translation"

### Supplementary Text

Supplementary material and methods

#### Cell culture

E14 mouse embryonic stem cells were obtained by the Tissue Engineering Unit of the Center for Genomic Regulation. Briefly, for maintenance culture cell were seeded onto plates pre-coated with 0,1% gelatin (ES-006-B, Millipore) under feeder-free conditions at a density of 1,5X$10^4$cells/$cm^2$ in knock-out DMEM supplemented with 15% embryonic stem cells qualified FBS (16141079, Gibco), 1% Penicillin/Streptomycin (15140122, Gibco), 1% L-Glutamine (25030081, Gibco), 1% Sodium Pyruvate (11360070, Gibco), 1% non-essential amino acids (NEAA) (11140068, Gibco) and Leukemia Inhibitory Factor (LIF) (ESG-1106, Millipore). For differentiation culture the medium was supplemented with 10%FBS. For normal maintenance, cells were passed any other day by dissociating them with the ACCUTASE® cell detachment solution (SCR005, Millipore). The ES Rex1-dGFP was obtained by Austin Smith's laboratory. For maintenance culture cells were plated at a density of 2 to 3X$10^4$cells/$cm^2$ always on pre-coated dishes. Cells were maintained in 2i medium made by mixing Neurobasal (21103048, Gibco) and DMEM/F12 (12634010, Gibco) 1:1 supplemented with N2 factor (17502-048, gibco) and B27 (17504-047, Gibco), 1% L-Glut, 1% NEAA, 1% Sodium Pyruvate, 1% Penicilin/Streptomycin, β-mercaptoethanol (31350010, Invitrogen), 1μM of PD0325901 (444968 Calbiochem) and 3μM of CHIR99021 (361571, Calbiochem). Cells were passed any other day. For lentiviral production, HEK293T packaging cell line was used. Cells were cultured in DMEM supplemented with 10% FBS and 1%L-Glutamine and 1% Penicillin/Streptomycin. All cell lines were pathogen tested.

#### Cell differentiation

To set up the Embryoid Body (EB) assay, E14 mESCs suspended in KO-DMEM supplemented with 10% serum at a density of 2x$10^4$cells/mL were seeded in hanging drops of 20μL onto the surface of a 150mm culture plate. After three days, the hanging drops containing the EBs were flushed and seeded on a low attachment plate. The medium was changed any other day until the day 10 of differentiation. In order to derive cardiomyocytes, at day three (D3) the cells were seeded on a gelatin pre-coated 96 multi-well plate and cultured for a maximum of 8 more days.

#### CRISPR/Cas9-mediated genome editing

Genome-editing was carried out by transfecting E14 cells with the bicistronic plasmid pSpCas9 PX459 (1μg per well in a 6-well plate) encoding for both the Cas9 and the sgRNAs targeting the first exon of either Sam68 or QKI. The transfected cells were then selected with puromycin and FACS sorted at clonal density in a 96 multi-well plate. Clones that were successfully edited were then Sanger-sequenced and the absence of the protein product verified by Western Blot. See Table S1 for gRNA sequences.

#### Western Blot

In order to extract proteins, an adequate volume of RIPA buffer (10mM Tris-HCl pH 8.0, 1mM EDTA, 1% Triton-X100, 0,1% sodium deoxycholate, 0,1% of sodium dodecilsulphate, 140mM of NaCl) supplemented with proteases inhibitor (1187358000111, Roche) was added to the cell pellet and kept on ice for 10 minutes pipetting at least once during the incubation. The cells were then centrifuged at max speed (>13'000 rpm) on a tabletop centrifuge for 10 minutes at +4°C. The supernatant containing the proteins is then preserved and the concentration of protein measured spectrophotometrically as absorbance at 595nm in Bradford reagent. An amount of proteins ranging from 25µg up to 60µg was then mixed with LDS 4X (NP007, Invitrogen) and loaded into a precast NUPAGE 4%-12% bis-tris gel (NP0321BOX, Invitrogen) immerged in MOPS buffer (NP0001, Invitrogen). The run was performed at 180V and the gel is dry-blotted with the iBlot™ 2 NC transfer stacks (IB23002, Invitrogen) and its corresponding device using the P3 program for 7 minutes. The nitrocellulose was then blocked with 5% milk in T-TBS (TBS buffer with 0,1% Tween20) for 1h at room temperature (RT). Afterwards this was incubated with the primary antibody in 2,5% milk in T-TBS for either 1h at RT or overnight at +4°C. Corresponding HRP-conjugated secondary antibody were then added in T-TBS with 2,5% milk and incubated for 1 hour at RT. To reveal the signal the membrane was incubated with the Immobilon Classico Western HRP substrate (WBLUC0100, Millipore) and revealed with the Amershan 600 (GE Healthcare and Life Sciences).

**Proliferation assay**

To assess the proliferation rate, $7{,}5X10^4$ cells were plated in a gelatin pre-coated 12 multi-well plate. Alive cells were then counted daily by Trypan Blue staining and quantified with the Countess™ II automated cell counter. Each well was counted at least twice. The graph in Figure 1B represents the average and the standard deviation of 3 independent experiments.

**Clonogenic assay**

For this assay, 200 cells were plated on a pre-coated 6 well plate and cultured with the E14 maintenance medium supplemented with LIF and 15% FBS. The cells were then left in culture for 10 days and the medium was changed every other day. At day 10 the colonies were stained with the Alkaline Phosphatase Staining Kit II (00-0055, Stemgent) according to the manufacturer's instructions. The colonies were then counted and the area of each colony quantified using Image J software. The dot plot in Figure 1D represents the average of 3 independent replicas.

**Lentiviral vectors production**

Lentiviral vectors were produced using the HEK293T packaging cell line. Briefly, 24 hours prior to transfection $9x10^6$ cells are plated in a 150mm diameter dish. A mix of 9µg of VSV-G envelope vector, 20µg of packaging **Δ**8.9 vector and 32µg of pLKO.1-shRNA mission vector (Sigma-Aldrich) was made and milli-Q water was then added up to a volume of 1125µl along with 125µl of 2,5M $CaCl_2$. The solution was incubated on a rotating wheel at RT for 5 minutes. Afterwards, 1250µl of 2XHBS pH 7.4 were added dropwise while vortexing at maximum speed. The resulting solution was incubated for 10 minutes at RT. Then this was added dropwise onto the cells. After 14-16 hours, the medium was changed and collected after 24 and 48 hours. The medium containing the virus was filtered through a 0,45µm filter and put into a polypropylene 25X36 cm tube (Beckman Culture). The viral particles were then concentrated through ultracentrifugation for 2 hours and 20 minutes at 22'000 rpm at 22 degrees Celsius. The pellet containing the virus was then resuspended in ice-cold PBS in order to concentrate the lentiviral vector 1000 times. The viruses were stored at -80°C

**Lentiviral infection and generation of Knock-down cell lines**

In order to quantify the exit from the pluripotent state we first generated the Knock-down and scramble Rex1-GFP cell lines via lentiviral spin infection. Briefly, for each infection were used 1,5X$10^5$ cell resuspended in 100µL of medium supplemented with polybrene at a concentration of 5 µg/mL. A variable amount (from 5 to 10µL) of lentiviral vector was used and the infection was performed by centrifugation at 1000G for 3 hours at 32°C. The cells were then resuspended and plated on a pre-coated 0,1% gelatin 6 well plate. The medium was changed 24 hours after plating and the selection of the successfully infected cells was performed with 10 µg/mL puromycin (P8833, Sigma-Aldrich) and started 48 hours after the infection.

**Exit from pluripotency assay**

The Rex1-GFP KD and scramble cell lines were used in order to assess the exit from pluripotency and the commitment of these cells. Briefly, the cells were allowed to differentiate by removing both LIF and the 2 inhibitors. Cells were then detached with the ACCUTASE® detachment solution and resuspended in ice-cold PBS supplemented with DAPI at a concentration of 0.1µg/mL in order to detect the fraction of death cells. Cells were analyzed at the BD LSR II cytometer. We first gated for forward and side scatter, we excluded the doublets and the death cells from the final analysis and we plotted the amount of residual GFP expression after 24, 48 and 72h after LIF and 2i removal. The bar plot in Figure 1F represents the average of 3 independent replicas.

**Teratoma assay**

All the experimental protocols were performed in accordance with the recommendations for the proper care and use of laboratory animals [local (law 32/2007); European (EU directive n° 86/609, EU decree 2001- 486) regulations, and the Standards for Use of Laboratory Animals n° A5388-01 (NIH)] and were approved by the local ethical committee (CEEA-PRBB). For Teratoma formation, 100µl of solution containing 500.000 mouse ES cells and 1:15 matrigel were injected into both flank sites of SCID BEIGE mice. The teratoma formation was stopped after 3 weeks of development. The resulting tissue was washed with PBS, fixed with 4%PFA, paraffin embedded, sectioned and stained with haematoxylin/eosin. Sections were evaluated and the presence of ectodermal, mesodermal and endodermal-derived tissue was quantified.
The teratoma quantification was performed by calculating the amount of each tissue in every teratoma slide analyzed. The different percentages of each experimental sample (the dots in the Figure S1D) show the average of three different portions of the same teratoma.

**RNA-sequencing**

In order to perform the RNA-sequencing, total RNA was extracted with the Maxwell 16 LEV simplyRNA Cells Kit (AS1270, Promega) according to manufacturer's instructions from WT, Sam68-/- and QKI -/- EBs in three biological replicates. DNA-free RNA was then quantified using the Nanodrop and quality checked with the Bioanalizer. rRNA depletion and library preparation was done with the TruSeq stranded total RNA Library prer Human/Mouse/Rat (20020596, Illumina). The sequencing was performed using 2x125bp paired-ends reads on a HiSeq 2500 sequencer with HiSeq v4 chemistry. The raw reads were de-multiplexed, by the bcl2fastq Illumina software. The Illumina Universal adaptor was trimmed off the raw reads with

Skewer (version 0.2.2), option "-x AGATCGGAAGAG". The quality of both raw and trimmed reads was assessed with the FastQC tool. Trimmed reads were aligned to the *Mus musculus* genome (mm10/GRCm38) with the STAR RNA-seq mapper (version 2.5.3a) using option "--quantMode GeneCounts" and the M14 version of the Gencode annotation in order to obtain raw counts per gene and per sample (the 4$^{th}$ column of the resulting counts file that corresponds to a "reverse stranded" protocol was kept). Bam files were converted to bigwig format with Samtools, so as to be loaded into the UCSC genome browser. The DESeq2 R/Bioconductor package was used to assess differential expression at gene level between experimental conditions. A gene was considered differentially expressed if the corresponding FDR-adjusted p-value was < 0.01 and the absolute fold change in log2 scale was > 0.6.
Last, we used vast-tools (version 2.0.0) in order to detect alternative splicing events. MATT was used to filter vast-tools combined output. In this case, a dPSI threshold of 25 (described as stringent by the authors) was applied to filter significant results.

**RNA retro-transcription, semi-quantitative and Real-Time PCR**

For RNA retro-transcription, a variable amount of RNA ranging from 500ng up to 1µg was retro-transcribed to cDNA using the NZY first-strand cDNA synthesis kit (MB125, NZYtech) according to manufacturer's instructions. Semi-quantitative Polymerase Chain Reaction (PCR) was performed by mixing the necessary amount of DNA with $H_2O$ up to a volume of 10µL. 10µL of NZYtaq II 2X green master mix (MB358, NZYtech) were then added. To amplify the DNA a specific thermocyclator program was used according to the melting temperature of the primers as well as to the specificity of each DNA target. The products of the PCR reaction were then run on an agarose gel (from 0,5 up to 3% depending on the amplimer size) in TAE buffer. The run was always performed at 100V and the gel analyzed with ChemiDoc XRS+ system by Biorad. Real-time PCR used for RNA-seq validation was performed by using SYBR® master mix (4309155, Applied Biosystem) according to manufacturer instruction. The reaction was run on a Viia™ 7 system and the fold enrichment derived with the ΔΔCt method.

**Ribosome profiling**

For each experiment of ribosome profiling we used 3 plates of 100mm of E14 cells at full confluence. Cells were treated with cycloheximide (01810, Sigma Aldrich) at a concentration of 100 µg/mL and were incubated for 15 minutes at 37°C. The media was withdrawn and the cells washed with ice cold PBS supplemented with cycloheximide, scraped and pelleted. The cells were then lysed with lysis buffer (20mM Tris-HCl pH7,4, 150mM of NaCl, 5mM of $MgCl_2$, 1mM DTT, 100 µg/mL, 1% of Triton and 25U/mL of Turbo DNase I [AM 2238, Invitrogen] in RNase free water) with the help of glass beads by vortexing 5 seconds for 3 times. The lysate was then left 10 minutes on ice and consequently centrifuged at 10'000 rpm for 10 minutes. To proceed we took an amount of RNA corresponding to 8-10 $OD_{260nm}$. In order to digest the unprotected RNA, RNAse I (AM2294, Invitrogen) at a concentration of 100U/µL was added and incubated at 25°C for 15 minutes. 10µL of SUPERase In (AM 2694, Invitrogen) at 20U/µL were added to stop RNase digestion and the sample loaded onto a sucrose cushion at 34% w/v in a polycarbonate ultracentrifuge tube (349622, Beckman Culture). The sample were then centrifuged at 70'000 rpm for 4 hours at +4°C. The supernatant was discarded and the pellet resuspended in the resuspension buffer of the Maxwell 16 LEV simplyRNA Cells Kit (AS1270, Promega). The RNA was then extracted according to the manufacturer instructions and resuspended in 40µl of RNAse free water. In order to select the fragments of interest we pre-run a 15% TBE-Urea polyacrylamide gel (EC6885BOX, Invitrogen) at 200V for 15 minutes in TBE buffer. 5µg of each sample were then mixed with the TBE-Urea sample buffer. The RNA was then denatured at 80°C for 90 seconds and loaded onto the gel and run at 200V for 65 minutes. The gel was then stained for 5 minutes with SYBR green (S32717, Invitrogen) and fragments between 28nt and 32nt excised. The bands were transferred into a RNase-free tube, resuspended in 360µl of RNase free water and the RNA extracted by crushing the gel with a pestle. Then it was incubated at 70°C for 10 minutes in order to melt the gel. Once melted the sample was loaded into a spin column with filter made of 2 pieces of whatmann paper and centrifuged at max speed for 2 minutes. Consequently, the RNA was precipitated by adding 40µL of 3M sodium acetate, 1µL of glycoblue (00585590, Invitrogen) and 500µL of isopropanol. The precipitation was performed either for 1 hour in dry ice or overnight at -80°C. Afterward the samples was centrifuged at 20'000g for 30 minutes at +4°C. The supernatant was discarded and the pellet washed with ethanol at 70% in RNase free water. It was then furtherly centrifuged at 20'000g for 2 minutes and then air dried for 10 minutes. Eventually, it was resuspended in 26µL of RNase free water. The ribosomal RNA was then removed by making use of the Illumina Ribo-zero rRNA removal kit (20020596, Illumina) according to manufacturer's instructions. The RNA left was then precipitated by adding 300µL of isopropanol 20µL of sodium acetate, 90µL of $H_2O$ and precipitated for 1 hour in dry ice. Then it was centrifuged again at 20'000g for 30 minutes at +4°C, the supernatant discarded and the pellet washed with 500µL of ethanol 70%, centrifuged again at 20'000g for 5 minutes at +4°C and the pellet was air-dried and resuspended in 10µL of TRIS-HCl pH8 10mM. 33µL of water were added and the samples were incubated at 80 °C for 90 seconds. In order to dephosphorylate the RNA, 5µL of T4 PNK buffer, 1µL of SUPERase IN and 1µL of T4 PNK enzyme (M0202L, New England Biolabs) were added and the mix incubated at +37°C for 1 hour. The enzyme was then inactivated by rising the temperature at up to +70°C. The RNA was then precipitated and washed as previously stated and resuspended in 8,5µL of RNase-free water. In order to add the linker, 1,5µL of pre-adenylated linker were previously denaturated (0.5µg/µL) at +80 °C for 90 seconds and then cooled down at RT. Then it was ligated by adding 2µL of T4 RNA truncated ligase buffer, 6µL of PEG 8000, 1µL of SUPERase In (20U/µL) and 1µL of T4 RNA truncated ligase enzyme (200U/µL, M024L, New England Biolabs), incubated for 2 hours and a half at RT. Then the RNA was precipitated and washed as previously described and resuspended in 10µL of Tris-HCl pH8 10mM. The samples were run on a 15% TBE gel as previously described, the RNA was excised, extracted and precipitated as previously described and resuspended in 10µL of TRIS HCl pH 8 10 mM. In order to convert it to cDNA 2µL of reverse transcription primer at 1.25µM were added along with 4µL of 5x first strand buffer, 1µL of dNTPs mix (10 mM), 1µL of 1M DTT, 1µL of SUPERase In (20 U/µL), 1µL of Superscript III enzyme (200 U/µL, 18080093, Invitrogen) and retro-transcribed at +50°C for 30 minutes. The cDNA was then precipitated and washed as previously described and resuspended in 10µL of water. The cDNA was then electrophoretically run, excised and precipitated as previously described. The cDNA was circularized adding 2µL of circ-ligase buffer, 1mM of ATP and 50mM of $MnCl_2$ and 100 units of CircLigase I (CL4111K, Epicentre). The sample was then incubated 1 hour at +60°C in a thermal cycler and then heat-inactivate the enzyme for 10 min at +80°C and the rRNA depleted once again. After precipitation and wash the cDNA was resuspended in 5µL of TRIS HCl pH8 10 mM and was PCR amplified in 100µL volume containing 20µL of Phusion HF buffer, 2µl of dNTPs mix (10 mM), 0.5µl of FW primer (100 µM), 0.5µl of RV barcoded primer (100 µM) and 71µL of RNase free water, 1µl of Phusion polymerase (2 U/µl, M05305, New England Biolabs). The PCR was performed with the following program: 1 cycle at 98°C for 30 sec, 14 cycles at 98°C for 10 sec, 65°C for 10 sec, +72°C for 5 sec. Eventually, the samples were run on an 8% (wt/vol) polyacrylamide non-denaturing gel for 40 min at 180V. The gel was then stained and a band at ~180nt was excised, extracted and purified as previously described. Then the libraries obtained were quality checked and sequenced using single-end 50bp sequencing on a HiSeq 2500 sequencer with HiSeq v4 chemistry. The bioinformatics analysis was performed by the bioinformatics core facility of the CRG. Briefly, the adaptor was trimmed off the raw sequences (fastq files) with skewer (version 0.2.2). The quality of both raw and trimmed reads was assessed with the FastQC tool. The reads that aligned (using bowtie2 version 3.2.0) to the rRNAs or the tRNAs (coordinates from the UCSC table browser), were removed (using bowtie2 with the following parameters: --no-unal --mp 3 -L 10 -N 1 --local -D 10 -R 3). All the remaining reads were size selected: only reads ranging from 22nt to 36nt were considered for further analysis in order to capture the ribosome protected fragments (approximately 30bp). The reads were then aligned to the Gencode M14 version of the genome (mm10/GRCm38) using the STAR mapper (version 2.5.3a) with parameters: --winAnchorMultimapNmax 100 --seedSearchStartLmax 15 –outFilterScoreMinOverLread 0 –outFilterMatchNminOverLread 0 –outFilterMatchNmin 10. Read coverage around the TSS was assessed, for each selected read length separated, in order to define an offset/shift value of ribosome position (the shifts are then applied to the read mapping position to obtain the approximate position of the translated codon). The reads were finally sorted out according to their alignment to either the coding sequence (CDS) or the 5' or 3' untranslated regions.

**Immunofluorescence**

EBs at D10 were collected and transferred into a 15ml conical tube. The medium was removed and the EBs fixed with 4% paraformaldehyde (PFA) for 30 minutes at room temperature. The PFA was removed and the EBs washed with PBS for 5 minutes. The EBs were then incubated for 30 minutes with solutions made of PBS with increasing concentration of sucrose (10, 20 and 30% in PBS). After that, the solution was carefully removed and the EBs were included in OCT and frozen. The tissue was then cut at the microtome and the put on a glass slide. In order to stain it, first the slide was put at +37°C for 30 minutes then it was washed 3 times in PBS. Afterwards, the slide was first incubated with blocking solution (5% BSA, 0,1% Triton-X100) and then with the primary antibody in a humid chamber overnight at +4°C. Then, the slide was washed 3 times for 5 minutes with PBS and incubated with a specific secondary antibody always in blocking solution for 1h at RT. Eventually, the slide was washed 3 more times in PBS for 5 minutes and it the slide was covered with a coverslip attached with the mounting solution containing DAPI to stain the nuclei. The image was acquired with a Leica TCS SP8 A0BS microscope at a magnification of 40X. The GFP intensity in Figure 4C was quantified with the ImageJ software by dividing the intensity of the GFP signal by the observed field-area and arbitrary choosing the value 1 to the WT sample.

**Computational analysis of the binding propensity**

The *cat*RAPID algorithm estimates the binding potential through van der Waals, hydrogen bonding and secondary structure propensities of both protein and RNA sequences allowing identification of binding partners with high confidence. As reported in a recent analysis of about half a million of experimentally validated interactions, the algorithm is able to separate

interacting vs non-interacting pairs with an area under the ROC curve of 0.78 (with False Discovery Rate FDR significantly below 0.25 when the Z-score values are > 2)
The specificity of the interaction of SAM68 or QKI for transcripts whose translation is affected in absence of Sam68 in differentiating EBs (Figure 4D) was calculated by considering the binding propensity multiplied by the RNA binding ability, as implemented in catRAPID omics v 2.0. The background distribution employed consisted of 1799 RBPs retrieved from literature. Similarly, the interaction of SAM68 or QKI with Gata4 (Figure 4E), was calculated by considering the binding propensity multiplied by the RNA binding ability, as implemented in catRAPID omics v 2.0 . To avoid biases arising from i) RNA type and ii) RNA length, we restricted the analysis of mouse transcriptome to protein coding transcripts that have length in the range (3000 to 4000 nts), i.e. similar to ENSMUST00000067417 (GATA4) which is 3400 nucleotides. The interaction binding profile (Figure 4F) was computed using the catRAPID fragment module.

#### Electromobility shift assay

Electrophoretic mobility shift assays were carried out using 10 fmol of radiolabeled probes incubated on ice for 15 min with His-$RBM_{Nter}$ in presence of 200 ng/µL yeast tRNA. Mixtures were then loaded on native 5% polyacrylamide gels, and migration was carried out in 0.5%TBE at 150V for 2 hours, followed by gel drying and Phosphorimager analyses.

#### Quantification and Statistical analyses

For each information about the statistical analysis, the meaning of the error bars and the number of replicates for each experiment please refer directly to the figure legends of each experiment.

### DATA AND CODE AVAILABILITY

#### Data and code availability statement

The datasets and code generated during this study are available as detailed in the Key Resources Table.

#### Data availability

All the data of the RNA-sequencing and the Ribosome Profiling are available at GEO with accession number GSE153800 and GSE153799 respectively

#### Computer code

The code used for analyzing and processing the PALM data can be found at https://gitlab.com/anders.sejr.hansen/palm_pipeline.
Please see Key Resources Table for a full list of all the codes and softwares and where to find them.

Alternative Splicing events for QKI
Ryr2(ex75) MmuEX0040863
Ttn (ex13) MmuEX0049851
Nebl (ex25) MmuEX0031345

Ttn (ex11) MmuEX0049849
Nebl (ex26) MmuEX0031346
Slmap (ex23) MmuEX0043713
Ank3 (ex24) MmuEX0004923
Sorbs2 (ex11) MmuEX0044378
Smpx (ex5) MmuEX0043997
Myo18a (Ex40) MmuEX0030455
Stim1 (ex10) MmuEX0045370
Slc8a1 (ex3) MmuEX0043591
Fhod3 (ex22) MmuEX0019238
Ttn (ex12) MmuEX0049850
Limch1 (ex9) MmuEX0026439
Kcnd3 (ex6) MmuEX0025030
Cacna1c (ex21) MmuEX0008749
Adgrg (ex6) MmuEX0021656

Alternative Splicing events for Sam68

Adgrg6 (ex6) MmuEX0021656
Mybpc3 (ex10) MmuEX0030286
Cacnac1c (ex22) MmuEX0008751
Cacnac1c (ex8) MmuEX0008750
Cacnac1c (ex30) MmuEX0008746
Cacnac1c (ex32) MmuEX0008748
Cacnac1c (ex21) MmuEX0008752
Cacnac1c (ex31) MmuEX0008747

qPCR primers

| | | |
|---|---|---|
| Actn2 | TGGCACCCAGATCGAGAAC | GTGGAACCGCATTTTTCCCC |
| Mybphl | CACTGGAGATAGCCTCATGCT | GTGCTCTTCAATAGAGGGCAG |
| Nebl | TACAACCCTCTGGGAAGTGC | GAGGGTGGCGTTTCCTTTAT |
| Myh6 | GCCCAGTACCTCCGAAAGTC | GCCTTAACATACTCCTCCTTGTC |
| Actc1 | CTGGATTCTGGCGATGGTGTA | CGGACAATTTCACGTTCAGCA |
| Tmod1 | TGAGCTAGATGAACTAGACCCTG | CGGTCCTTAAATTCCTTCGCTTG |
| Tnni3 | TCTGCCAACTACCGAGCCTAT | CTCTTCTGCCTCTCGTTCCAT |
| Tnnc1 | GCGGTAGAACAGTTGACAGAG | CCAGCTCCTTGGTGCTGAT |
| Tnnt3 | GGAACGCCAGAACAGATTGG | TGGAGGACAGAGCCTTTTTCTT |
| Scn5a | ATGGCAAACTTCCTGTTACCTC | CCACGGGCTTGTTTTTCAGC |
| Ryr2 | TCAAACCACGAACACATTGAGG | AGGCGGTAAAACATGATGTCAG |
| Rbm20 | GGCCAAAACAAGCCCGATATT | CCCTGTCTGAGGTAGGCTCT |
| Rbm24 | TTTTGCCTTTGGCGTTCAACA | GCTGCACATGGGGAATGAC |
| Gata4 | CCCTACCCAGCCTACATGG | ACATATCGAGATTGGGGTGTCT |
| Gata5 | | |
| Mef2c | ATCCCGATGCAGACGATTCAG | AACAGCACACAATCTTTGCCT |
| Nkx2-5 | GACGTAGCCTGGTGTCTCG | GTGTGGAATCCGTCGAAAGT |

| | | |
|---|---|---|
| Tbx18 | GTACCTGGCTTGGCACGAC | GCATTGCTGGAAACATGCG |
| Isl1 | CAGTCCCAGAGTCATCCGAGT | TGGGTTAGCAGTTTTGTCGTT |
| Tnnt2 | CAGAGGAGGCCAACGTAGAAG | CTCCATCGGGGATCTTGGGT |
| Myl2 | ATCGACAAGAATGACCTAAGGGA | ATTTTTCACGTTCACTCGTCCT |
| Cacnb2 | ACTAGAGAACATGAGGCTACAGC | GCACTATGTCACCCAAACTGGAT |
| Lama1 | CAGCGCCAATGCTACCTGT | GGATTCGTACTGTTACCGTCACA |
| 18s | GTAACCCGTTGAACCCCATT | CCATCCAATCGGTAGTAGCG |
| Rplp0 | TTCATTGTGGGAGCAGAG | CAGCAGTTTCTCCAGAGC |

Alternative splicing primers

| | | |
|---|---|---|
| Ryr2 | CCTCAGACCCAGAGAGGACAG | AGTAGTTTGTGCCACACAGCT |
| Titin | GCAACCAAAGCCAAAGAGCAA | GCTGCTCTGGGACCTTTGTG |
| Stim1 | CTCTCAACATCGACCCCAGCT | TCTGAGATCCCAGGCCAAGC |

RNA-Immunoprecipitation primers

| | | |
|---|---|---|
| Gata4.2 | CACCCCAATCTCGATATGTTTGA | GCACAGGTAGTGTCCCGTC |
| Gata4.3 | CACGCTGTGGCGTCGTAAT | CTGGTTTGAATCCCCTCCTTC |

| REAGENT or RESOURCE | SOURCE | IDENTIFIER |
|---|---|---|
| Antibodies | | |
| Anti-Sam68 C-20 | Santa Cruz | Sc-333 |
| Anti-quaking N-20 | Santa Cruz | Sc-514468 |
| Mouse anti-beta tubulin | Abcam | Ab6046 |
| Normal Rabbit IgG | Santa Cruz | Sc-3888 |
| Mouse Anti-Gata4 | CRG protein facility | |
| Alexafluor488 donkey anti-mouse | Life Technologies | A21202 |
| Alexafluor555 goat anti-rabbit | Life Technologies | A32732 |
| Protein-G-HRP conjugated | Abcam | Ab97046 |
| Goat anti-rabbit-HRP conjugated | Dako | P0448 |
| Anti-Sam68 H4 | Santa Cruz | sc-514468 |
| Anti-panQKI N147/6 | Millipore | MABN624 |
| Chemicals, Peptides, and Recombinant Proteins | | |
| Nocodazole | Nocodazole | M1404 |
| Leukemia inhibitory factor | ESGRO | ESG1106 |
| Cycloheximide | Sigma-Aldrich | 239763 |
| N2 | Gibco | 17502-048 |
| B27 | Gibco | 17504-047 |
| Critical Commercial Assays | | |
| Alkaline phosphatase staining kit II | Stemgent | 00-0055 |
| Maxwell 16 LEV simplyRNA Cell Kit | Promega | AS1270 |
| TrueSeq stranded total RNA Library prer Human/Mouse/Rat | Illumina | 20020596 |
| Ribozero rRNA removal kit | Illumina | 20020596 |
| Oligonucleotides | | |

| | | |
|---|---|---|
| sgRNA targeting Sam68 GGCGCGAGGCAGGATCGTCC | This paper | n/a |
| sgRNA targeting Quaking GGTCGGGGAAATGGAAACGA | This paper | n/a |
| Preadenylated linker for Ribomose profiling libraries 1/5rApp/CTGTAGGCACCATCAAT/3ddC/ | This paper | |
| Reverse transcription primer for Ribosome profiling libraries 5′(Phos)AGATCGGAAGAGCGTCGTGTAGGGAAAG AGTGTAGATCTCGGTGGTCGC(SpC18)CACTCA(S pC18)TTCAGACGTGTGCTCTTCCGATCTATTGAT GG TGCCTACAG-3′ | Ingolia et al., 2012 | PMID: 22836135 |
| Forward primerfor Ribosome profiling library 5′-AATGATACGGCGACCACCGAG ATCTACAC-3′ | Ingolia et al., 2012 | PMID: 22836135 |
| Reverse barcoded primer for Ribosome profiling libraries 5′CAAGCAGAAGACGGCATACGAGATNN NNNNGTGACTGGAGTTCAGACGTGTGCTCTTCC G-3′ | Ingolia et al., 2012 | PMID: 22836135 |
| Primers for qPCR see table S1 | This paper | n/a |
| Primers for semi-quantitave PCR see table S2 | This paper | n/a |
| Recombinant DNA | | |
| pLKO.1-shRNA mission vector bacteria Sam68 | Sigma-Aldrich | |
| pLKO.1-shRNA mission vector bacteria QKI | Sigma-Aldrich | |
| Software and Algorithms | | |
| *cat*Rapid | Livi et al., 2016 | http://s.tartaglialab.com/page/catrapid_group |
| Bowtie2 | Langmead and Salzberg, 2012 | http://bowtie-bio.sourceforge.net/bowtie2/index.shtml |
| Samtools | Li et al., 2009 | http://samtools.sourceforge.net/ |
| Skewer | Jiang et al., 2015 | doi: 10.1186/1471-2105-15-182. |
| Bioconductor | Huber et al., 2015 | doi: 10.1038/nmeth.3252. |
| DESeq2 | Love et al., 2014 | doi: 10.1186/s13059-014-0550-8. |
| Bwa-mem | Li et al., 2009 | [PMID: 19451168] |
| Vast-tools | Tapial et al., 2017 | *Genome Res*, 27(10):1759-1768 |
| MATT | Gohr et al., 2019 | https://doi.org/10.1093/bioinformatics/bty606 |

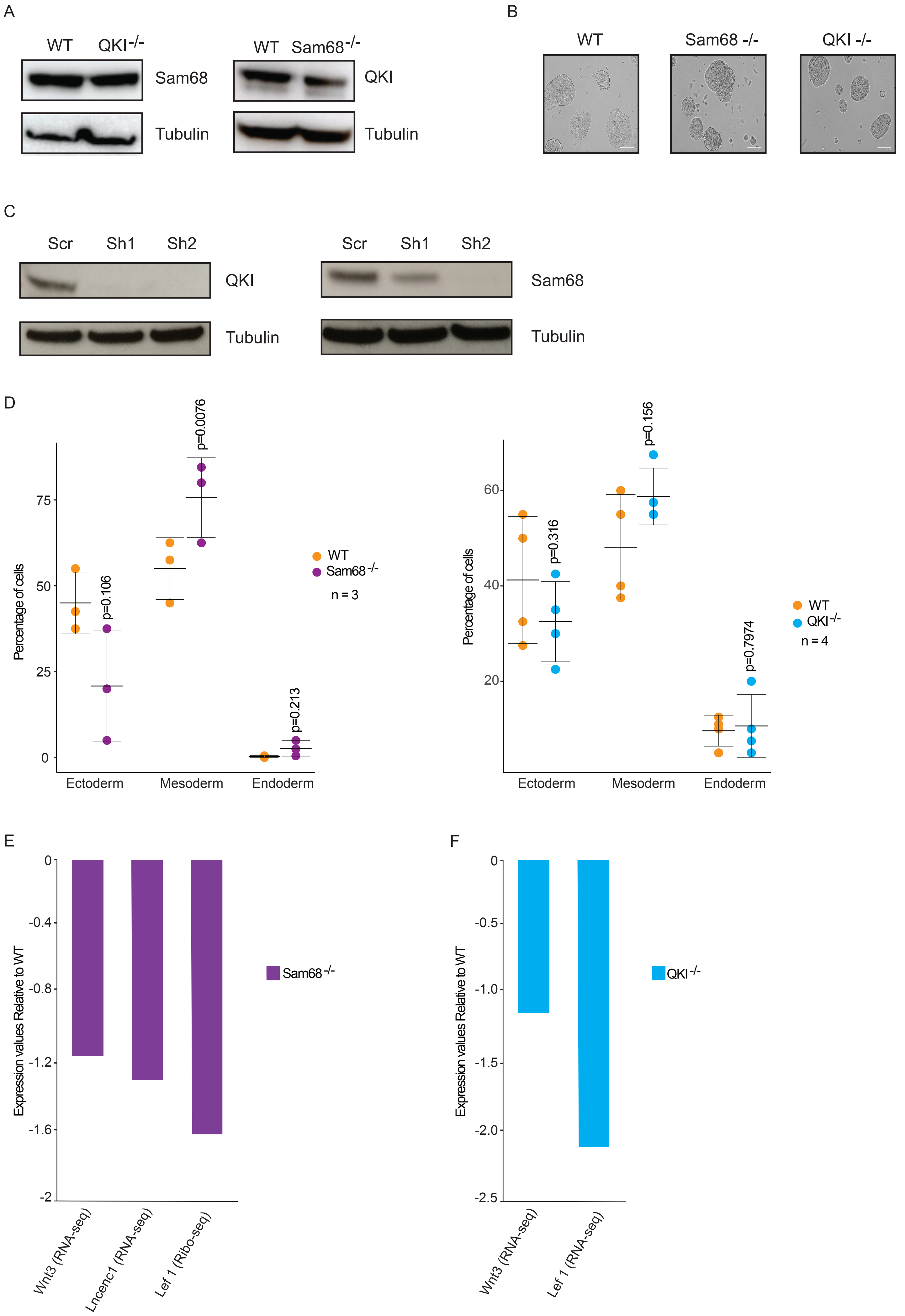
A
WT QKI-/-
Sam68
Tubulin
WT Sam68-/-
QKI
Tubulin
B
WT
Sam68 -/-
QKI -/-
C
Scr Sh1 Sh2
QKI
Tubulin
Scr Sh1 Sh2
Sam68
Tubulin
D
Percentage of cells
p=0.106
p=0.0076
p=0.213
WT
Sam68-/-
n = 3
Ectoderm Mesoderm Endoderm
p=0.316
p=0.156
p=0.7974
WT
QKI-/-
n = 4
Ectoderm Mesoderm Endoderm
E
Expression values Relative to WT
Sam68-/-
Wnt3 (RNA-seq)
Lncenc1 (RNA-seq)
Lef 1 (Ribo-seq)
F
Expression values Relative to WT
QKI-/-
Wnt3 (RNA-seq)
Lef 1 (RNA-seq)

Supplementary figure1

a) WB showing absence of compensation at the protein level of one member of the family in the KO background of the other one

b) KOs and WT mESCs colonies in bright-field microscopy

c) WB showing the KD protein levels of either QKI or Sam68 in Rex1-dGFP cells

d) Quantification of the teratoma derived tissues obtained by injecting WT and either Sam68$^{-/-}$ (on the left) or QKI$^{-/-}$ (on the right) mESCs in SCID mice

e) mRNA expression and RNA translation efficiency of mESCs pro-proliferative factors in Sam68$^{-/-}$ mESCs compared to the WT control

f) mRNA expression and RNA translation efficiency of mESCs pro-proliferative factors in QKI$^{-/-}$ mESCs compared to the WT control

A

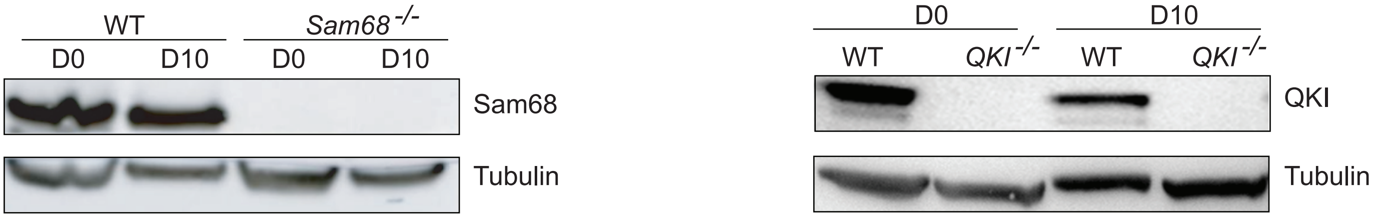

B

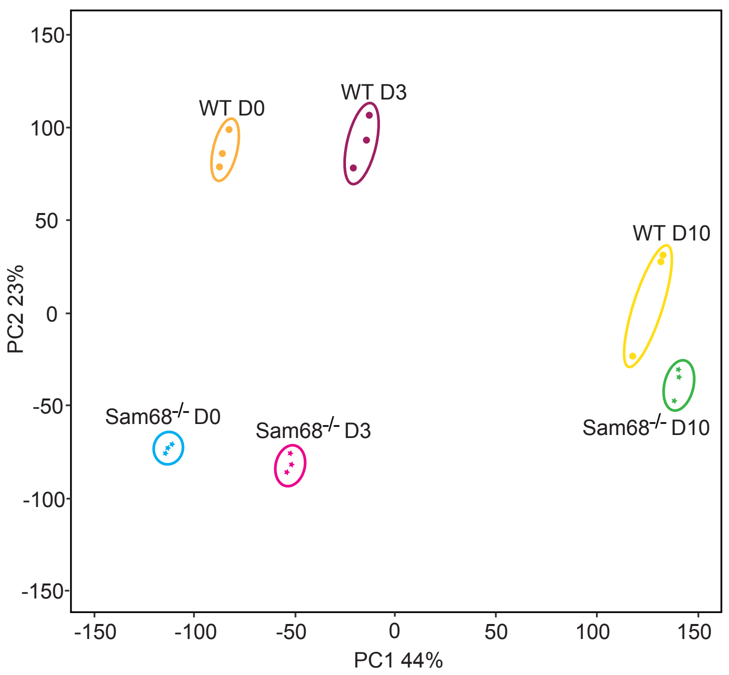

C

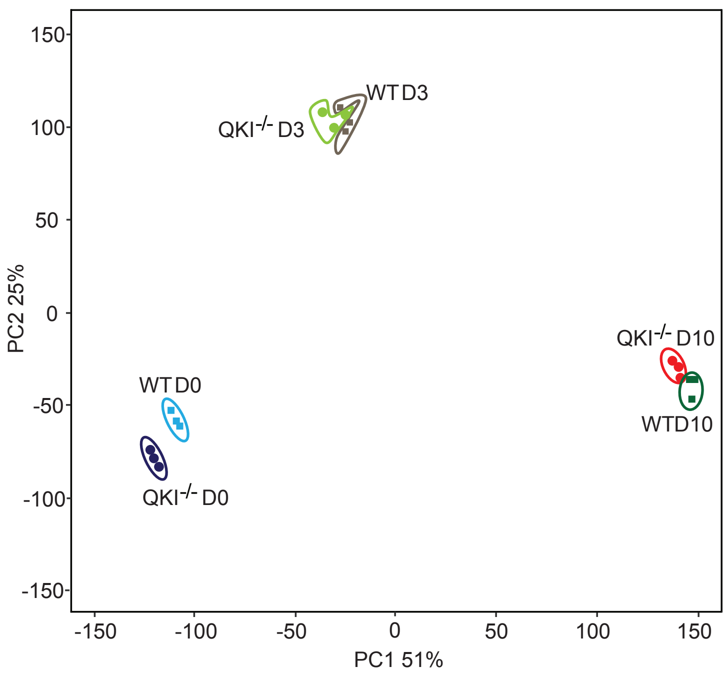

D

| | | Protein Coding | lnRNAs | miRNAs | Others |
|---|---|---|---|---|---|
| QKI | D0 | 1158 | 242 | 155 | 3456 |
| | D3 | 218 | 104 | 7 | 1569 |
| | D10 | 941 | 19 | 0 | 46 |
| Sam68 | D0 | 2501 | 51 | 19 | 112 |
| | D3 | 2867 | 5 | 7 | 10 |
| | D10 | 3647 | 14 | 1 | 49 |

Supplementary figure 2

a) WB showing the expression levels of Sam68 in WT and Sam68 KO undifferentiated mESCs and EBs at day 10 of differentiation

b) Principal component analysis of the RNA-sequencing WT and Sam68$^{-/-}$ samples

c) Principal component analysis of the RNA-sequencing WT and QKI$^{-/-}$ samples

d) Table showing the absolute numbers of differentially expressed RNAs for each RNA category. Only RNAs with ± 1,5 fold change and with an adjusted p value<0,01 are represented

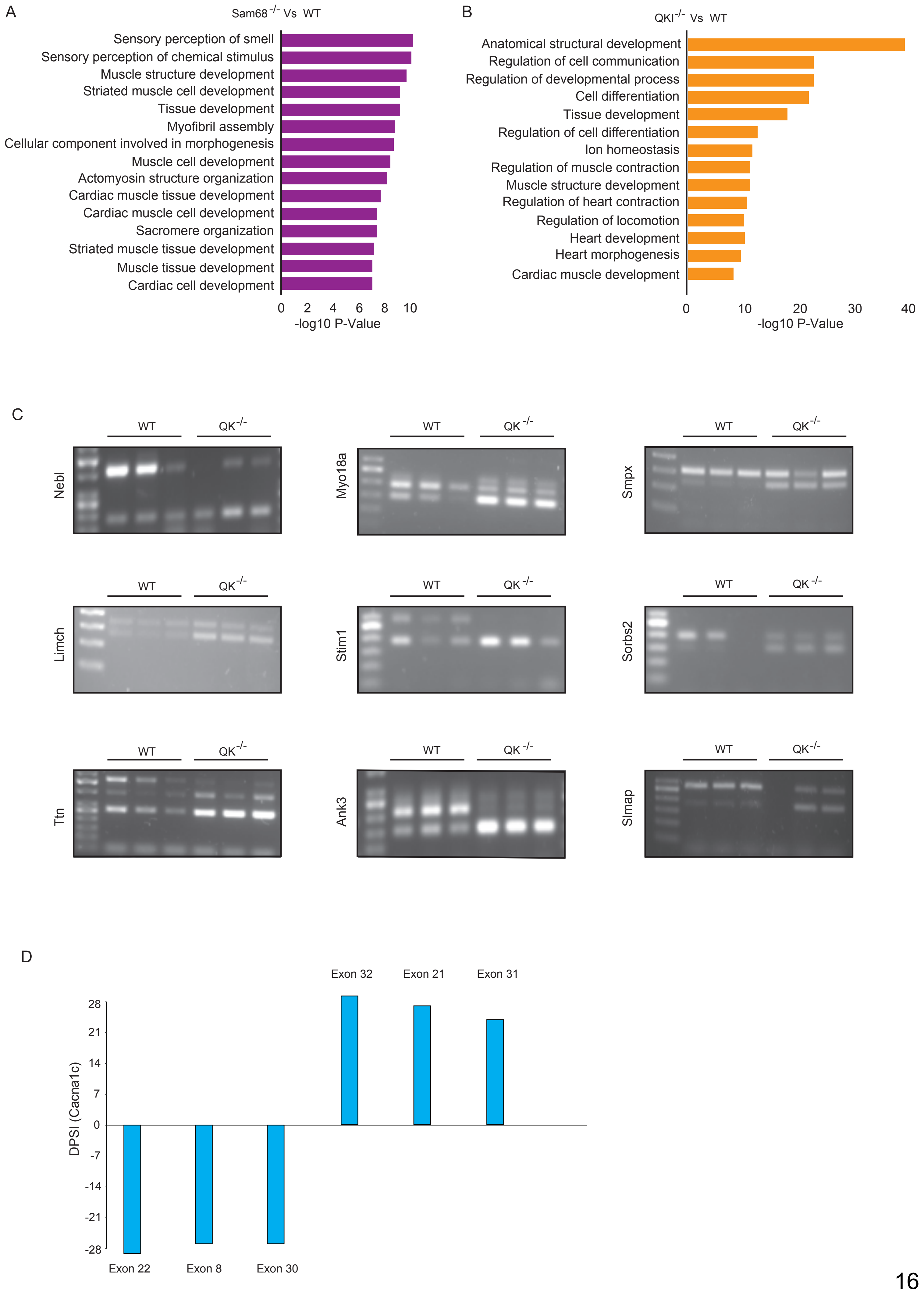
A
Sam68 -/- Vs WT
Sensory perception of smell
Sensory perception of chemical stimulus
Muscle structure development
Striated muscle cell development
Tissue development
Myofibril assembly
Cellular component involved in morphogenesis
Muscle cell development
Actomyosin structure organization
Cardiac muscle tissue development
Cardiac muscle cell development
Sacromere organization
Striated muscle tissue development
Muscle tissue development
Cardiac cell development
0 2 4 6 8 10
-log10 P-Value
B
QKI-/- Vs WT
Anatomical structural development
Regulation of cell communication
Regulation of developmental process
Cell differentiation
Tissue development
Regulation of cell differentiation
Ion homeostasis
Regulation of muscle contraction
Muscle structure development
Regulation of heart contraction
Regulation of locomotion
Heart development
Heart morphogenesis
Cardiac muscle development
0 10 20 30 40
-log10 P-Value
C
WT QK -/-
Nebl
Myo18a
Smpx
Limch
Stim1
Sorbs2
Ttn
Ank3
Slmap
D
DPSI (Cacna1c)
28 21 14 7 0 -7 -14 -21 -28
Exon 22
Exon 8
Exon 30
Exon 32
Exon 21
Exon 31

Supplementary figure 3

a) Gene ontology analysis of the most upregulated genes in Sam68 KO embryoid bodies at day 10 of differentiation shows overrepresentation of cardiac-related terms

b) Gene ontology analysis of the RNA-sequencing of the most downregulated genes in QKI KO embryoid bodies at day 10 of differentiation shows overrepresentation of cardiac- related terms

c) Validation of aberrant cassette-exons AS on cardiac-related RNAs in $QKI^{-/-}$ EBs at day 10 of differentiation

d) Bar plot showing several aberrant AS cassette-exon events occurring on the Cacna1c transcript in $Sam68^{-/-}$ EBs at day 10 of differentiation

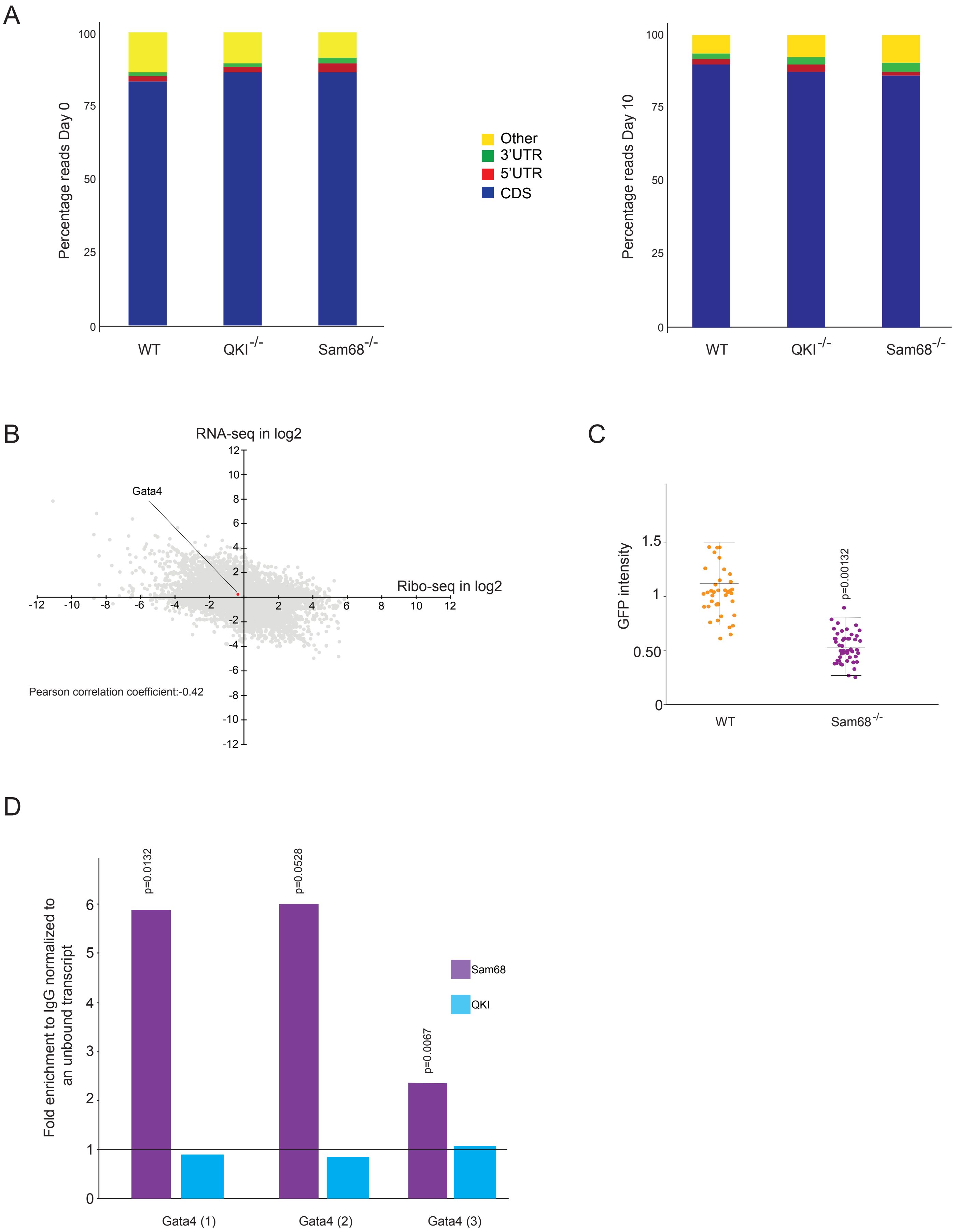
A
Percentage reads Day 0
100
75
50
25
0
WT
QKI-/-
Sam68-/-
Other
3'UTR
5'UTR
CDS
Percentage reads Day 10
B
RNA-seq in log2
Ribo-seq in log2
Gata4
Pearson correlation coefficient:-0.42
C
GFP intensity
1.5
1
0.50
0
p=0.00132
WT
Sam68-/-
D
Fold enrichment to IgG normalized to an unbound transcript
p=0.0132
p=0.0528
p=0.0067
Sam68
QKI
Gata4 (1)
Gata4 (2)
Gata4 (3)

Supplementary figure 4

a) Bar plot showing that the majority of the ribosome profiling reads at day0 (left) and day10 (right) of EBs differentiation align to the codifying sequence of the RNAs demonstrating the good quality of the generated libraries

b) Plot that shows the correlation between transcription and translation changes in QKI KO EBs at day 10 of differentiation

c) Quantification of immunofluorescence shown in figure 4C

d) RNA immunoprecipitation of Sam68 and QKI to detect Gata4 binding. The bar plot shows

the real-time PCR demonstrating the enrichment of the Gata4 transcript compared to IgG and normalized to a non-bound RNA (Rplp0).
